# Spatially-specific working memory activity in the human superior colliculus

**DOI:** 10.1101/2020.06.17.157669

**Authors:** Masih Rahmati, Kevin DeSimone, Clayton E. Curtis, Kartik K. Sreenivasan

## Abstract

Theoretically, working memory (WM) representations are encoded by population activity of neurons with distributed tuning across the stored feature. Here, we leverage computational neuroimaging approaches to map the topographic organization of human superior colliculus (SC) and model how population activity in SC encodes WM representations. We first modeled receptive field properties of voxels in SC, deriving a detailed topographic organization resembling that of the primate SC. Neural activity within male and female human SC persisted throughout a retention interval of several types of modified memory-guided saccade tasks. Assuming an underlying neural architecture of the SC based on its retinotopic organization, we used an encoding model to show that the pattern of activity in human SC represents locations stored in WM. Our tasks and models allowed us to dissociate the locations of visual targets and the motor metrics of memory-guided saccades from the spatial locations stored in WM, thus confirming that human SC represents true WM information. These data have several important implications. They add the SC to a growing number of cortical and subcortical brain areas that form distributed networks supporting WM functions. Moreover, they specify a clear neural mechanism by which topographically organized SC encodes WM representations.

**Significance Statement:** Using computational neuroimaging approaches, we mapped the topographic organization of human superior colliculus (SC) and modeled how population activity in SC encodes working memory (WM) representations, rather than simpler visual or motor properties that have been traditionally associated with the laminar maps in the primate SC. Together, these data both position the human SC into a distributed network of brain areas supporting WM and elucidate the neural mechanisms by which the SC supports WM.

## INTRODUCTION

We have known for decades that working memory (WM), a critical building block for higher cognition, relies on the persistent activity of neurons selective for the memoranda (Funahashi et al., 1989; Miller et al., 1996). More recent theoretical work suggests that WM features are encoded by the joint activity of large numbers of neurons with tunings that span the memorized feature space (Ma et al., 2014). Evidence from human studies using functional magnetic resonance imaging (fMRI), which allows for the simultaneous measurement of brain activity at scales that tile the space of neural population, supports the notion that macro-level distributed patterns of activity encode WM representations (Christophel et al., 2012; Riggall and Postle, 2012; Sreenivasan et al., 2014; Ester et al., 2015). Computational model-based methods designed to identify these information-containing patterns demonstrate that WM representations are present in a surprising number of cortical brain regions (Sreenivasan and D’Esposito, 2019). The success of these methods depend both on how encoding of the feature is distributed across a neural population (e.g., how visual space is distributed over V1) and how precisely aggregate voxel-wise measures of neural activity are feature-tuned (Brouwer and Heeger, 2009; Naselaris et al., 2011). It follows that WM representations of visual space are robustly encoded in areas with systematic retinotopic organization, like early visual cortex (Sprague et al., 2014; Rahmati et al., 2018), and to lesser extents in areas with coarse topographic organization, like frontal and parietal cortex (Jerde et al., 2012; Mackey et al., 2017).

In this study, we focus on a subcortical structure, the human superior colliculus (SC), and test its potential role in spatial WM. We are motivated by the growing appreciation for the important role that subcortical regions play in cognition (Basso and May, 2017; Halassa and Kastner, 2017), the well-established topography of the cat and macaque SC (Cynader and Berman, 1972; Goldberg and Wurtz, 1972a), and the above-mentioned recent developments in modeling fMRI measures of population activity. Traditionally, the SC mediates orienting behaviors (e.g., gaze shifts) by coordinating activity between two tightly registered eye-centered topographic maps: a map representing input from the visual system and a map representing motor output in the form of the angle and amplitude of saccades (Wurtz and Albano, 1980; Sparks, 2002; Gandhi and Katnani, 2011). However, two recent lines of evidence argue against a strictly visuomotor-centric model of SC function. First, pharmacological inactivation of the macaque SC motor map induces a form of visual neglect akin to extinction, but does not cause anopsia (i.e., visual field defects) or paresis (i.e., deficits in voluntary oculomotion) (McPeek and Keller, 2004; Lovejoy and Krauzlis, 2010; Zénon and Krauzlis, 2012). Second, neural activity in the cat and macaque SC motor map encodes the spatiotopic locations of a behavioral goal, rather than the specific metrics of the saccade to the goal (Freedman and Sparks, 1997; Bergeron et al., 2003). Therefore, the functional role of the SC cannot be explained simply in terms of visual input or motor output. Instead, the SC may integrate signals computed throughout the brain into a common topographical organization, acting as a staging area for organizing flexible and goal-oriented behavior into a map of space that is weighted by bottom-up salience and top-down goal-relevance (Fecteau and Munoz, 2006). Indeed, studies that use memory-guided saccade (MGS) tasks to dissociate visual and saccade related activity often observe delay period activity in macaque SC (Shen et al., 2011; Sadeh et al., 2018), consistent with the notion that the SC plays a role in spatial WM.

Here, we leverage computational neuroimaging approaches to 1) map the topographic organization of human SC, and 2) assuming an underlying neural architecture based on this topography, model how population activity in SC encodes WM representations disentangled from simple visual or motor components.

## Materials and Methods

### Participants

Six subjects (ages 27**-**49; 5 male; one left-handed) participated in the study. The subjects were in good health with no history of psychiatric or neurological disorders, had normal or corrected-to-normal visual acuity, and gave informed written consent. The study was approved by the New York University Committee on Activities Involving Human Subjects and the New York University Abu Dhabi Institutional Review Board.

### Stimulus display

We generated stimuli and interfaced with the MRI scanner, button-box, and eye-tracker using MATLAB software (The MathWorks, Natick, MA) and Psychophysics Toolbox 3 (Brainard, 1997; Pelli, 1997). Stimuli were presented using a PROPixx DLP LED projector (VPixx, Saint-Bruno, QC, Canada) located outside the scanner room and projected through a waveguide and onto a translucent screen located at the end of the scanner bore. Subjects viewed the screen at a total viewing distance of 64 cm through a mirror attached to the head coil. The display subtended approximately 32° of visual angle horizontally and vertically. A trigger pulse from the scanner synchronized the onsets of stimulus presentation and image acquisition.

### Eye tracking

To ensure fixation compliance and to record saccadic responses, we measured eye gaze constantly throughout the experiment using a MRI compatible Eyelink 2K (SR Research). Using the freely available iEye toolbox (github.com/clayspacelab/iEye), we preprocessed and scored eye-tracking data automatically, quantified the precision (average standard deviation of tangential and radial components of the saccade landing points) and response times of visual and memory guided saccades, and plotted example time-courses and trajectories of saccades shown in Figure 2C and D. Subjects were able to reliably fixate through the delay; fixation breaks (saccade amplitude > 2° from fixation) during the delay only occurred in 0.5% to 2.0% of the trials in five out of six subjects. The remaining subject made quick saccades away from and then back to fixation on 7.0% of trials. Given the relative infrequency of unwanted saccades, we did not exclude any trials from the fMRI analyses.

**Fig. 1.**
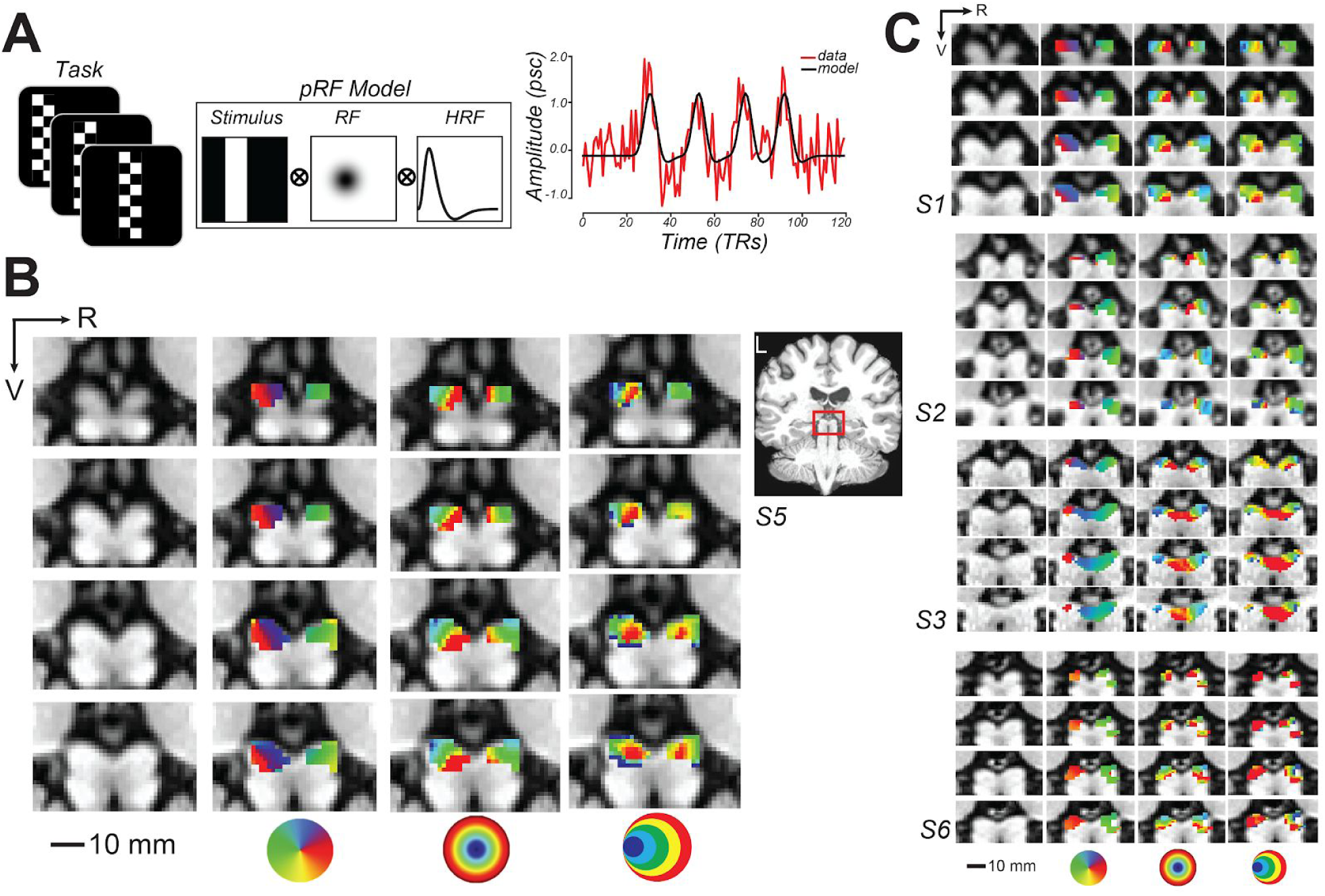
Topographic mapping of human superior colliculus (SC). ***A***. To model voxel population receptive fields (pRF), subjects viewed bars of contrast reversing checkerboards that swept across the visual field. Bar positions over time converted into binary apertures were projected onto a 2D Gaussian model of a receptive field (RF) and convolved with a hemodynamic response function (HRF). To the right, a single sample voxel in SC is plotted for one run. ***B***. Enlarged coronal slices through the human SC in an example subject (red box inset). R = right; V = ventral. From left-to-right, the columns depict the T1 anatomy, polar angle, eccentricity, and size parameter maps of an example subject (S5; thresholded at *r*^*2*^ ≥ 0.1). The colored circles are visual field keys. ***C***. Topography of SC is consistent across other subjects.

**Fig. 2.**
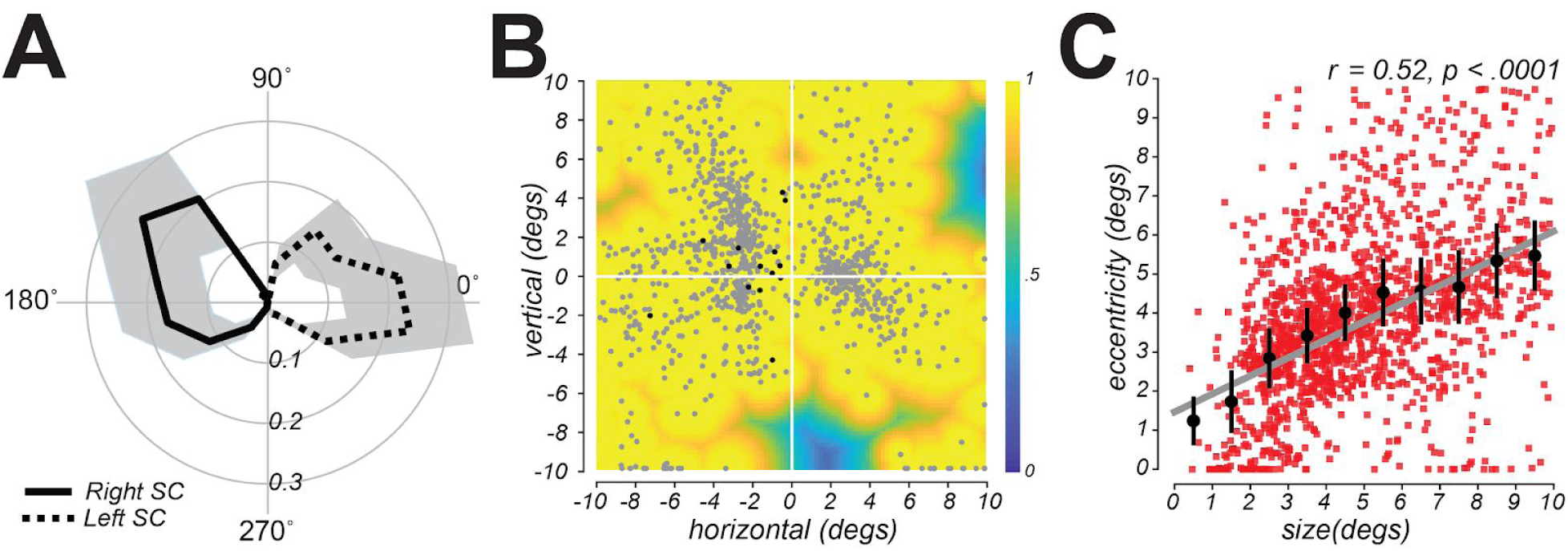
*A*. Radial histograms of pRF polar angle in SC demonstrate strong contralateral coverage of the visual field. Based on pRFs from all subjects, the lines are the mean fractional volume representing each polar angle (± SEM). ***B***. Aggregate field of view (FOV) when pRF location and size parameters are combined. Each gray dot represents the center of single voxel pRFs. The color represents the maximum pRF value across the population of voxels in the SC and reflects the relative effectiveness of visual stimulation in evoking a response in the SC. Black dots (n=14) denote pRFs from the left SC with a center in the ipsilateral left visual field (no such ipsilateral centers were found in the right SC). ***C***. Size of voxel pRFs in the SC increased linearly with eccentricity. Red squares = SC voxels from all subjects; Black dots = binned means (± SEM); Gray line = linear fit.

### MRI data acquisition

MRI data were acquired in the Center for Brain Imaging at NYU with a 3-Tesla Siemens Prisma MRI scanner scanner using a 32-channel head coil. Twenty functional series of 120 volumes were collected for the retinotopic mapping task and twenty functional runs (except for one subject from whom we collected ten runs) of 232 volumes were collected for the SWM task. Each functional run was acquired with 14 coronal slices and a gradient echo, echo planar sequence with a 128 square matrix, 192 mm field of view, and 2.0 mm slice thickness, leading to a voxel size of 1.5 ⨯ 1.5 ⨯ 2.0 mm (TR = 1.5 s, TE = 41 ms, flip angle = 66°, bandwidth = 752 Hz/pixel). A partial Fourier factor of 7/8 was used to acquire an asymmetric fraction of *k*-space and GRAPPA parallel imaging with a factor of 2 was used to reduce acquisition time. The posterior edge of the acquisition volume was aligned in the mid-sagittal plane with the posterior edge of inferior colliculus. We also collected a high-resolution T_1_-weighted MPRAGE (0.8 mm isotropic voxels, 256 ⨯ 240 mm) in order to register functional scans to an anatomical image. In addition, for each scanning session we collected a single whole-brain-coverage functional image (TR = 10.8 s) with the same spatial resolution as the partial-brain coverage functional images in order to align the partial-coverage functional images to the whole-brain anatomical images.

### Population receptive field (pRF) mapping

We used established procedures to model the pRF parameters in SC voxels (Dumoulin and Wandell, 2008; DeSimone et al., 2015). During scanning, subjects were presented with a checkerboard-patterned bar whose elements reversed contrast with a full-cycle frequency of 8 Hz (Fig. 1A). The bar subtended 8° of visual angle across its width and extended beyond the boundaries of the screen along its length. The bar was oriented either vertically or horizontally and swept across the screen perpendicular to the bar orientation, passing through central fixation. Each scanning run consisted of four 30 s sweeps (left to right, right to left, top to bottom, and bottom to top) in a random order, with 12 s mean-luminance blank periods at the start and end of the run. Subjects performed a demanding fixation task that required them to detect and map the color of the fixation cross (which could turn red, green, blue, or yellow every 1.5 s) to one of four button presses. We motion-corrected and co-registered functional data with the anatomical images. For each voxel, we removed the linear trend and converted the time-series to z-units. For the mapping experiment, we modeled each voxel in terms of a Gaussian pRF (Fig. 1A) using methods and tools previously described (DeSimone et al., 2015, 2016). The pRF model provides a description of each voxel’s BOLD response in terms of a retinotopic location and extent. We also modeled the delay of the hemodynamic response function (HRF) and the baseline of the BOLD signal. The delay parameter estimates the time to peak and time to undershoot of the HRF. The baseline parameter ensures that the modeled and measured BOLD signals vary about a single global mean. In an initial phase of the parameter estimation, we used a sparse and coarse grid search with an effective stimulus down-sampled by 2D bilinear interpolation to 5% of the original resolution. The best fit from the sparse sampling of model parameter space was then used as a seed in the final phase of a fine-tuned gradient-descent error minimization using the non-resampled stimulus. For each subject the SC region-of-interest was drawn based on anatomy and a pRF model threshold of *r*^*2*^ ≥ 0.1. Note that the pRF model failed in one subject (S4) even when lowering the cutoff threshold and we could not discern topography in SC. Thus, for this subject we selected all voxels within the SC based on anatomic T1 images for further analysis. Importantly, our spatial WM results were not dependent upon subject S4; in fact, the results were statistically more robust when excluding S4, although we include S4 in the results presented below for completeness. Using procedures similar to (Winawer et al., 2010), we estimated the field of view (FOV) of the SC map from the full pRF model. To represent the FOV of the full SC map in visual space, we used 2D Gaussians whose positions within the visual field and widths were determined by each voxel’s pRF center and size parameters, and whose maximum value equalled 1. We did this on the pRF parameters aggregated across the left and right SC of all subjects. Since many points in the visual field were covered by several pRFs, when combining the pRFs we mapped each visual field coordinate to the maximum pRF value.

### Spatial working memory experiments

In order to measure spatial WM representations in the SC, we imaged the brain while subjects maintained a location in WM during a long memory retention interval (Fig. 3A). Trials began with a brief visual stimulus (full contrast circle with the radius of 0.25 visual angles) presented for 300 ms in the periphery at one of eight angular locations evenly spaced from 22.5 to 337.5° of polar angle in 45° intervals and jittered by ±10°, at an eccentricity of 9-11° of visual angle (Fig. 3B). The color of the visual stimulus indicated the transformation required to remap its location to the goal of a later memory-guided saccade (MGS). A white stimulus indicated no transformation; green indicated a MGS to the location mirrored across the horizontal meridian; red indicated a MGS to the location mirrored across the vertical meridian; and blue indicated a MGS to the location mirrored across both the horizontal and vertical meridians. After a 10.5 s memory delay, a black dot appeared for 400 ms at a random uniformly sampled polar angle (0-360°) and radius (9-11° of visual angle) from central fixation. Subjects first made a visually-guided saccade (VGS) to this target, and then immediately made a MGS to the transformed location guided by memory (1.4 s alloted for both saccades). Finally, the stimulus was re-presented at the correct transformed location as a means of providing performance feedback (500 ms). After subjects made a corrective saccade to the feedback target, an inter-trial interval (ITI) followed of 9.8 s of central fixation that preceded the next trial. Each scanning run contained 16 trials (each 22.5 s), sampling each of the 8 angular locations twice. Each of the four MGS transformation conditions were counterbalanced across pairs of successive scanning runs. All subjects practiced outside of the scanner before the experiment began. Two task manipulations were designed to help us understand the nature of the information maintained in the SC during the memory delay. First, the intermediate VGS prevented subjects from being able to plan, and potentially maintain, the metrics of the MGS during the delay. Second, the various transformations moved the task-relevant location — the goal of the MGS — to a position in visual space that was independent of the retinal position of the visual stimulus. Together, we aimed to eliminate simply visual and/or motor components while honing in on WM representations of visual space.

**Fig. 3.**
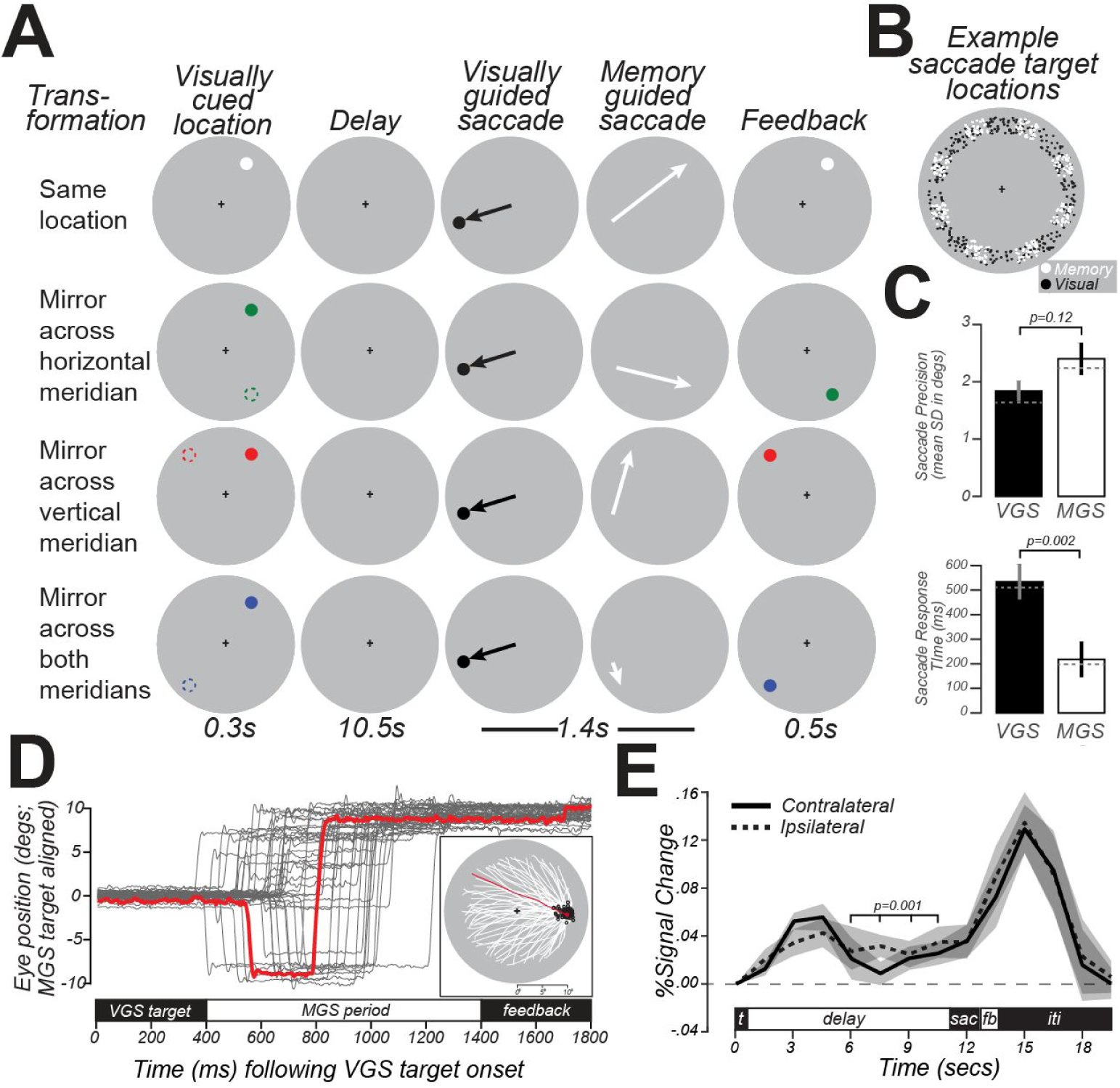
Task schematic, behavioral data, and delay-period activity in human SC. ***A***. Schematic of four types of MGS trials. In each condition, trials began with a brief visual target located in the periphery (colored dots; left column). Following a delay, subjects made a VGS to a target whose location was unpredictable. Then, subjects immediately made a MGS to a location based on the initial visual target. In one condition, the MGS was directed to the visual target. In the other conditions, the MGS was made to simple geometric transformations of the visual target (dashed circles, left column; for reference here but not displayed). These included mirror transformations across each meridian and both meridians. The color of the visually-cued target indicated the type of transformation. Feedback was provided after the MGS with a visual stimulus at the correct location. Because of the VGS, the metrics of the MGS could not be predicted. The transformations dissociated the goal of the MGS from the visually-stimulated retinal position. ***B***. Locations of VGS and MGS targets were distributed 9-11° in the periphery. ***C***. VGS were slightly more accurate and significantly slower than MGS. Bars = mean (± SEM); Gray dashed line = median. ***D***. Example eye-tracking traces (gray lines) from a subject during one scanner session. All trials are rotated such that despite the various transformations of the visual target location, all MGS targets are rotated to a location 10° to the right. The red trace highlights an example trial. In the inset, we re-plot only the MGS trajectories (white lines), which start from a wide variety of peripheral locations following the VGS but converge and end (black circles) near the aligned MGS target location. The red trace highlights an example trial. ***E***. Group-averaged (± SEM) BOLD signal in SC voxels persisted significantly above pre-trial baseline (gray dashed line) during the delay period for trials when the MGS target was in the contralateral and ipsilateral hemifield. The delay was defined as the average of the last four TRs in the delay period, identified by the bracket above the time courses.

### Inverted Encoding Model

To reconstruct a representation of SWM from the pattern of SC activity during the memory delay, we used a spatial inverted encoding model (IEM; Fig. 4) (Brouwer and Heeger, 2009). First, we modeled each voxel’s response as a weighted sum of eight information channels, each in the form of a squared one-dimensional cosine function centered at one of the nine equally spaced polar angles around an invisible ring. We estimated voxel-channel weights by fitting a general linear model to a subset of data used only for training. For this training, we only used trials in which the visual stimulus and MGS target were co-located (“Same location” condition, Fig. 3A). We then inverted these regression weights to estimate the contribution of each channel to a representation of visual-space in the held-out data from the other conditions that required a spatial transformation of the visual stimulus. Finally, we averaged all information channels, weighted by their estimated channel contribution, to reconstruct the population’s representation. For visualization purposes, we depict the information channels arranged around an invisible ring centered in the visual field. We estimated the population activity in each trial by averaging each voxel’s BOLD activity during the last four TRs of the delay period. To increase the signal-to-noise ratio, we combined trials by computing a two-fold mean trial time-series, reducing the total number of trials by half while maintaining the counterbalancing of the exemplars across the memory locations. We repeated the IEM training and reconstruction procedure using a 10,000 iteration bootstrap procedure with different arrangements of trials for computing the two-fold mean time-series. This ensured that any effects were not simply due to bias in the sampling and recombination of trials.

**Fig. 4.**
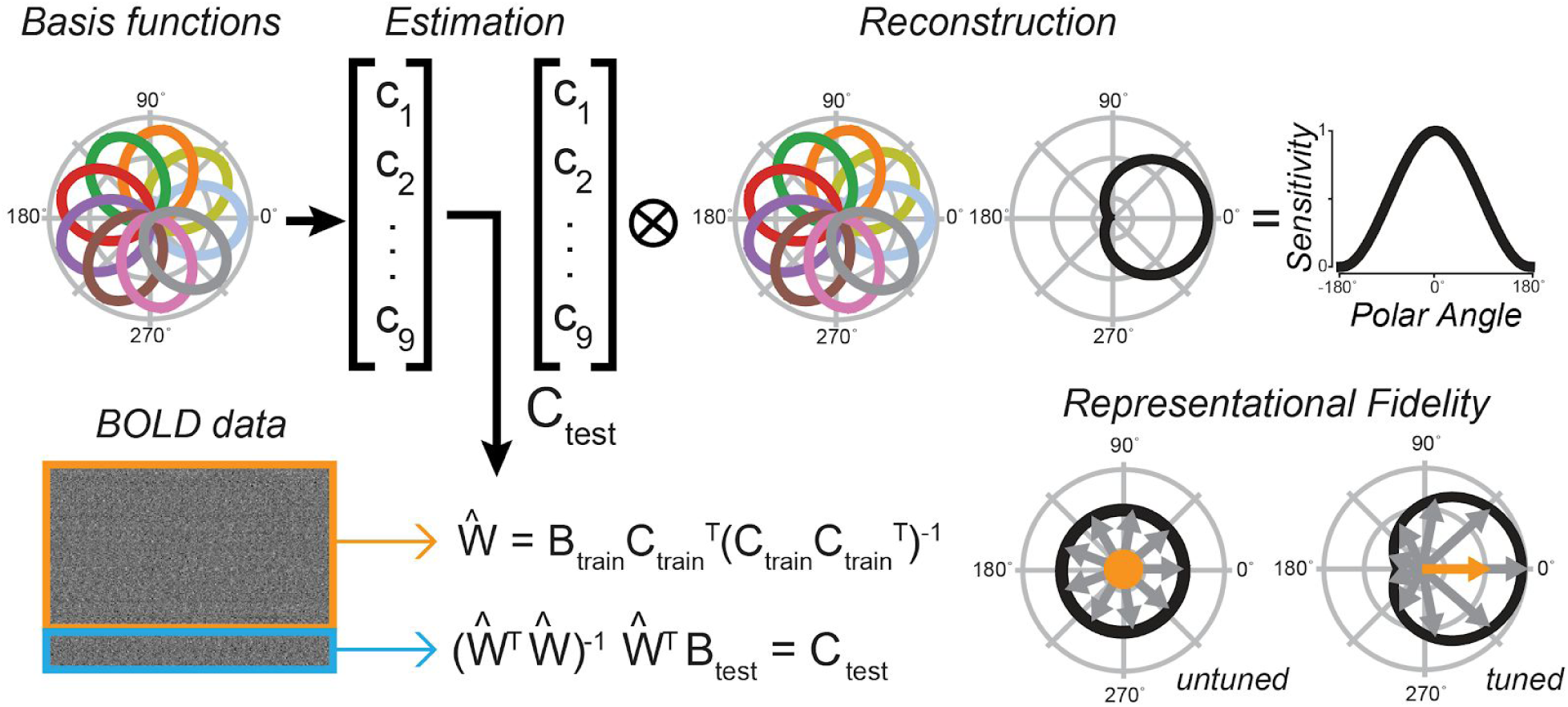
Inverted encoding model (IEM). Using a standard IEM, we calculated regression weights (W) from a training set of BOLD data (B_train_; orange box) and corresponding hypothetical channel coefficients (C_train_) represented by nine evenly spaced radial basis functions, each tuned to a specific angle. We calculated the contribution of each basis function in the final reconstruction (C_test_) by linearly combining a new set of BOLD data (B_test_; blue box) and the inverse of the regression weights. To reconstruct a representation of visual space, we used a linear combination of all basis functions, each weighted by its corresponding contribution in C_test_. To the right, we unwrap the curve to show a sample sensitivity profile across angles in visual space. We calculated representational fidelity — a metric for the goodness of reconstructions — as the vector mean of a set of unit vectors around different angles, each weighted by the reconstructed sensitivity at that position. Displayed are examples of poor/untuned, and good/tuned representations. Conceptually, our model provides a means to map a multivoxel population response into the coordinates of visual space.

### Statistical analysis

To quantify the goodness of our reconstructions, we used a modified version of the representational fidelity metric (Sprague et al., 2016). Representational fidelity quantifies the similarity between a given reconstruction and a standard tuning function; however, this can be overly sensitive to the gain of the reconstruction peak at the cost of sensitivity to deviations from the reconstruction center. To adjust the sensitivity of the fidelity metric, we included a cost function in our modified fidelity calculation (Eq. 1), where *f*_*stndrd*_ and *f*_*recon*_ are the standard and reconstructed tunings (both circular), *l* is the location (0 - 360°), *g* is the cost function, *θ*_*f_stndrd*_ is the parameters set of the standard tuning function, and *err*_*recon*_ is the deviation(error) of the reconstruction peak from the true location.

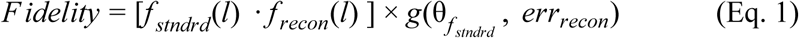

To validate the significance of our reconstructions, we built 10,000 IEMs, each trained after shuffling the training data, and compared the fidelity distributions corresponding to the real and permuted data through a nonparametric Kolmogorov-Smirnov test at the individual subject level and a paired t-test, after a 1000 iteration bootstrap across subjects, at the group level.

In order to link the pRF model of retinotopy and the spatial IEM, we compared each voxel’s polar angle preference derived from the two models. For the pRF model, we simply used the polar angle of the pRF center. For the IEM, we summed all information channels weighted by their estimated regression coefficients yielding a polar angle tuning curve for each voxel. Since the IEM estimates were derived from the task in which all stimuli were 9-11° in the periphery, we restricted our analysis to SC voxels whose pRF centers were at least 5° in eccentricity. We then calculated the circular correlation coefficient between the pRF polar angle and the peak of the IEM tuning curve.

### Data availability

Analysis code and experimental data are available at https://osf.io/mkgd2/.

## Results

### Retinotopic mapping

Following our pRF mapping procedures (Fig. 1A), we examined the modeled receptive field properties of voxels in the SC by overlaying the pRF model parameters on the T1 anatomical image (shown for a representative subject in Figure 1B and individual subjects in Figure 1C). We found orthogonal polar angle and eccentricity representations of the visual field along the SC. The topography revealed a graded upper-to-lower visual field representation along the medial-to-lateral axis of the SC, and a graded foveal-to-peripheral visual field representation along the anterior-to-posterior axis. The SC pRFs largely tiled the contralateral visual field (Fig. 2A). The “bow-tie” shape of the distribution of polar angles, as found in previous fMRI studies of the retinotopy of the SC (Schneider and Kastner, 2005; DeSimone et al., 2015), seems to imply a relative underrepresentation of angles near the vertical meridian. However, when we consider the full receptive field model and combine the pRF centers and sizes to estimate the FOV of the SC, it is clear that visual stimulation at most retinal locations effectively drives the SC (Fig. 2B). Moreover, like in the macaque (Marino et al., 2008), we found a positive correlation between the size and eccentricity of pRF parameters in the SC (Pearson’s *r* = 0.52, *p* < 0.0001; Fig. 2C). Overall, our model of the topographic structure of the human SC closely resembles that of the cat and macaque (Cynader and Berman, 1972; Goldberg and Wurtz, 1972a) and a previous report in humans (DeSimone et al., 2015).

### Spatial working memory

The latency and precision of both visual and memory-guided saccades were similar to previous studies that used delayed saccade or antisaccade tasks (Curtis and Connolly, 2008; Saber et al., 2015), despite the transformations and double saccades (Fig. 3C/D). The average BOLD signal in the retinotopically defined human SC persisted above pre-trial baseline during the memory period suggesting it may play an important role in WM (percent signal change; mean = 0.04, 95% CI = [0.009 0.08], *p* < 0.01; Fig. 3E). However, it was not lateralized with respect to the MGS target - a pattern incongruous with the clear lateralization of the SC pRFs (Figs. 1 and 2). This may be due to the intervening VGS target and complex transformations required by the task. Alternatively, averaging over many voxels may result in a measure that is too coarse to capture the population dynamics by which the SC encodes WM.

Motivated by our pRF findings, we used a multivoxel model of visual space (Fig. 4) to test whether topographic patterns of activity in human SC encode WM locations. Conceptually, the model provides a means to map a multivoxel population response into the coordinates of visual space. We assumed an underlying neural architecture based on the retinotopic organization of the voxels within SC, and modeled each voxel’s response with a set of basis functions that tiled polar angle space. We tested this assumption by comparing the angles derived from the pRF model with the angles derived from the IEM for each voxel in the SC. First, we trained the model using trials in which no transformation of the visually-cued target was required, deriving modeled basis functions from which we estimated each voxel’s preferred polar angle. Second, using circular correlation, we confirmed that the pRF and IEM polar angle parameters were similar (*r* = 0.26, *p* < 0.002). This suggests that the two forward-modeling approaches converged on very similar polar angle parameters despite the differences in modeling (pRF and IEM) and differences in cognitive demands (visuospatial attention and WM). Next, using the IEM-based model trained on no transformation trials, we tested it on trials requiring transformations (see *Materials and Methods*). Consistent with the notion that SC population delay activity encodes spatial information in WM, our model could accurately reconstruct the transformed location of the MGS (Fig. 5A, right). Importantly, these locations stored in WM were computed from spatial transformations of the visual targets and thus were not locations that were retinally stimulated earlier in the trial. Models trained on the location of the visual target or the VGS location were unable to reconstruct these locations (Fig. 5A, left and center), indicating that SC delay activity encoded the abstract representation of the memory location rather than the visually presented targets. Quantification of these results using our modified representational fidelity metric confirmed that SC population activity during the delay was spatially tuned only for the location of the MGS target (mean fidelity = 0.0013, 95% CI = [0.0010 .0023], *p* < 0.001; Fig. 5B). Remarkably, we found these effects at the individual level in every subject except S4, the subject in which we could not discern topography in the SC based on our pRF model (Fig. 5C). Overall, the results were consistent and provide robust evidence for spatial WM encoding in topographically organized human SC.

**Fig 5.**
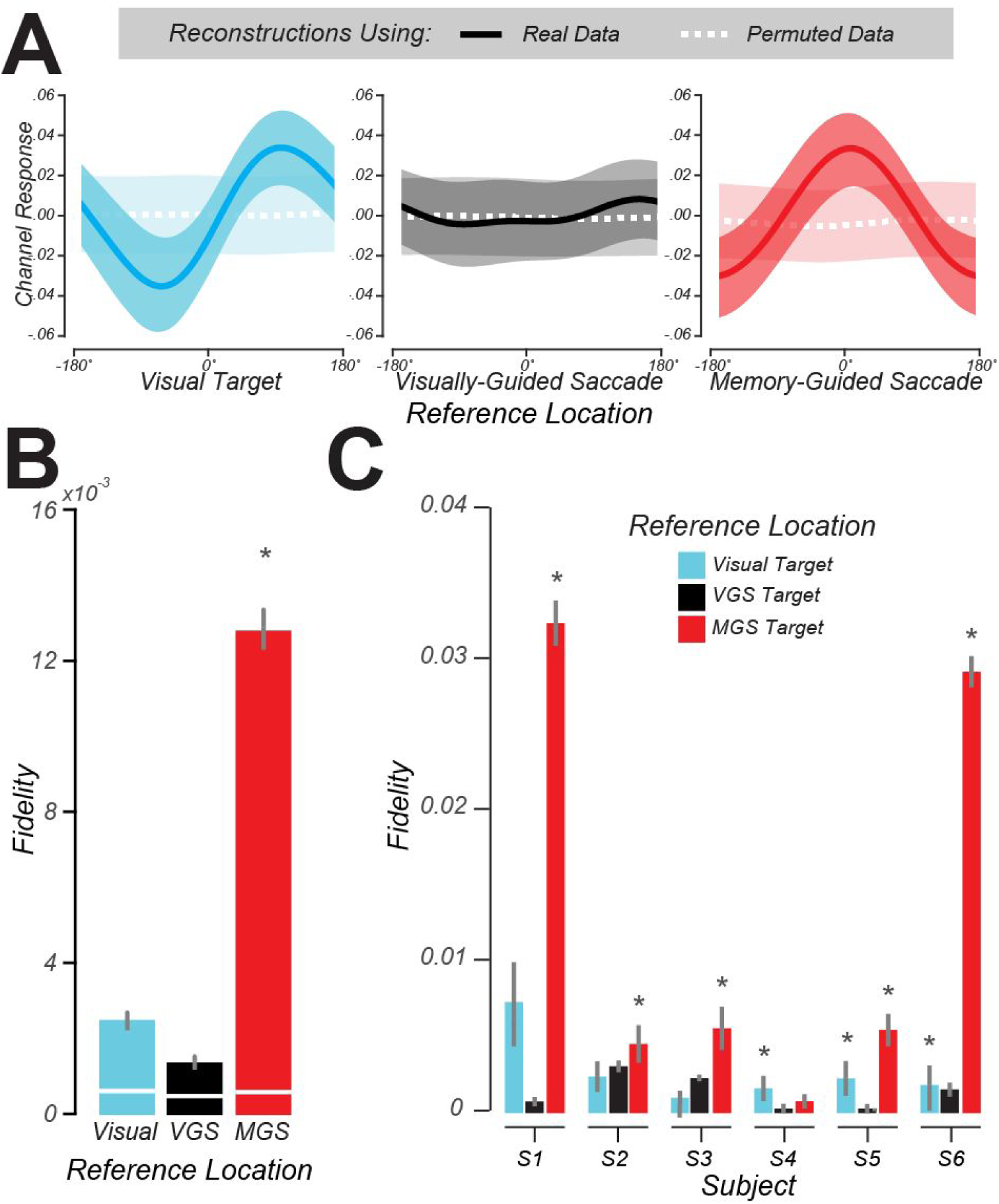
Modeling WM representations in human SC. ***A***. We used delay period activity in human SC to reconstruct visual space. From left to right: the average reconstructed sensitivity (± SEM) in visual space aligned to the visually-cued target, VGS target, and MGS target locations, respectively. In each panel, all trials are aligned to the corresponding reference location centered at 0°. The dashed white lines depict reconstructions from BOLD data with the trial labels permuted. ***B***. Representational fidelity (± SEM) corresponding to three reference locations, compared to shuffled data (white lines) computed at the group level. Note that the SC population activity during the delay is largely tuned for the visual-spatial location of the MGS (*p* < 10^−18^), not the visual target or VGS. ***C***. Even at the individual subject level, we find greater fidelity for the MGS location for all subjects except the one subject whose pRF model failed (S4). In three subjects there was smaller tuning for the visually-cued target, but this small effect was not significant at the group level.

## Discussion

Motivated by the increasing role that the subcortex (Halassa and Kastner, 2017) and specifically the SC (Basso and May, 2017) is thought to play in cognition, we utilized recent developments in fMRI modeling to test how WM representations are encoded in the population activity of the human SC. The topographic organization of the SC can be leveraged in models of how populations encode WM representations (Serences and Saproo, 2012). Using model-based fMRI, we showed that the topography of human SC resembles that of macaque SC, the activity in retinotopic SC persisted during WM retention intervals, and at the population level it encoded the spatial location of WM representations.

### Retinotopy of human SC

Using pRF mapping, we identified a visual field map in human SC that systematically represented the contralateral visual field. The structure of the map corresponded to the topographic organization of the cat and macaque SC (Cynader and Berman, 1972; Goldberg and Wurtz, 1972a) in the following ways. The representations of upper and lower portions of visual fields were found in the medial and lateral, respectively, parts of the SC map. The representations of foveal and peripheral portions of the visual field were found in the anterior and posterior, respectively, parts of the SC map. Similar to that reported in human visual cortex (Dumoulin and Wandell, 2008), we found that the size of the estimated receptive field of voxels correlated with its eccentricity, where smaller receptive fields were nearer the foveal. Both of these observations matched those reported using electrophysiology in macaques (Goldberg and Wurtz, 1972a; Marino et al., 2008). Previous fMRI studies using phase-encoding methods for retinotopic mapping (Schneider and Kastner, 2005, 2009; Katyal et al., 2010) found similar results with respect to the orderly contralateral maps of polar angle. Comparable polar angle maps in human SC have also been derived during the generation of saccades of different angles (Savjani et al., 2018), suggesting that mapping evoked by saccades and visual stimulation are in registration or perhaps originate from the same map. These previous studies also reported the same anisotropic distribution of angles we observed, with a lesser representation along the upper and lower vertical meridians compared to the horizontal meridian. The cause and significance of the anisotropy, which is also ubiquitous in cortical retinotopic areas, remains a point of debate (Larsson and Heeger, 2006; Winawer et al., 2010; DeSimone et al., 2015). However, by using the full receptive field model to combine the pRF centers and sizes we were able to estimate the FOV of the SC, which clearly covers the whole visual field. Our pRF methods enabled us to extend characterizations of the human SC by mapping its eccentricity axis along the rostral-caudal axis of the SC, mapping the receptive field size of its voxels, and using the full model to estimate the FOV of the SC.

### Spatial working memory

During the WM delay period, BOLD activity in SC was low but clearly persisted above pre-trial baseline throughout the delay. On the one hand, this appears to align well with electrophysiological recordings from macaque SC neurons that typically show a slow but increased rate of discharge prior to saccades, including memory-guided saccades (Paré and Wurtz, 2001; Shen et al., 2011). On the other hand, the delay activity was not contralateralized with respect to the location of WM targets as would be expected based on the discharge properties of macaque SC neurons (Robinson, 1972) and the contralateral organization of the SC map in our subjects. Of course, our task’s visuomotor transformations could disrupt a lateralized response. But perhaps our coarse averaging of BOLD signals across all voxels from each side of the SC was affected by the distributed neural activity related to foveating the central fixation stimulus (Krauzlis et al., 2017) and/or inhibitory neural activity related to suppressing unwanted saccades to the remembered location during the delay (Ikeda et al., 2015). Although the delay activity suggests that human SC plays some role in spatial WM - an important finding in itself - it does not tell us how it might contribute to WM.

Therefore, motivated by the spatiotopic organization of the SC, we constructed a multivoxel model of how population activity in SC encodes spatial WM representations. Similar encoding models of fMRI data have been useful in testing hypotheses about how *cortical* areas store relevant features of WM representations (Ester et al., 2015; Rahmati et al., 2018; Cai et al., 2019). Our results demonstrate that these models also work well in the subcortex, as we were able to model the population response in the SC that encoded spatial WM representations. Critically, the patterns of delay period activity we modeled did not encode retinal positions of past visual stimuli or future planned saccades. The locations held in WM were abstract transformations of visually stimulated locations, and the visually-guided saccades negated strategies involving the maintenance of saccade motor metrics. Therefore, the pattern of activity across the human SC neural population encodes abstract, cognitively defined locations in the absence of visual stimulation or motor commands.

The SC encoding of abstract locations in WM may be initiated by feedback signals from the cortex. If so, we might ask: what is the nature of these signals and from where do they originate? With respect to their nature, they resemble the spatial attention effects that have been described in the macaque SC (Krauzlis et al., 2013). Visually-evoked SC responses are larger when the stimulus is behaviorally relevant and the goal of a saccade (Goldberg and Wurtz, 1972b). When task related saccades are dissociated from the locus of attention, SC neurons with RF matching an attended target also show enhanced discharge rates (Ignashchenkova et al., 2004) and manipulations using microstimulation and chemical inactivation provide causal support for the role of the macaque SC in covert attention (Cavanaugh et al., 2006; Lovejoy and Krauzlis, 2010). Attention also causes enhanced neural responses measured with fMRI in the human SC (Schneider and Kastner, 2009; Katyal et al., 2010; Katyal and Ress, 2013). Indeed, one of the key mechanisms supporting spatial WM may involve sustained covert attention (Awh and Jonides, 2001; Jerde et al., 2012). In the context of our results, therefore, attention related signals targeting neurons with receptive fields matching the transformed locations may sculpt the population encoded responses in the SC we observed. Likely sources of these top-down influences include brain areas with direct connections to the macaque SC, namely, lateral prefrontal cortex, frontal eye field (FEF), lateral intraparietal (LIP) area, and V1, each of which are known to support spatial WM functions (Sommer and Wurtz, 2001; Armstrong et al., 2009; Koval et al., 2011; Everling and Johnston, 2013; van Kerkoerle et al., 2017). Moreover, human fMRI studies employing encoding models like the one used here consistently report that spatial WM representations are encoded in the patterns of population activity in these same cortical regions (Jerde et al., 2012; Riggall and Postle, 2012; Ester et al., 2015; Rahmati et al., 2018).

However, for a number of reasons it is unlikely that SC simply integrates cortical commands and relays them to brainstem oculomotor plants. In the macaque, representations of visual priority emerge more rapidly in SC than in V1, indicating that feedback signals from SC may sculpt the gain of responses in cortex (White et al., 2017). The SC has more ascending projections through the pulvinar and mediodorsal thalamus that could influence cortex, for example, than descending projections arriving into the SC (May, 2006). Moreover, lesions to the SC impair behaviors that depend on covert attention, but surprisingly do not affect the typical attentional enhancement of neuronal activity in extrastriate cortex (Zénon and Krauzlis, 2012). Therefore, the SC may play critical roles in spatial cognitions like attention and WM through circuits that both interact with but at the same time are independent of the cortex.

Our results depended on a model of the well-defined spatial topography of the SC, where space is systematically distributed over hundreds of voxels, and an assumption that WM was encoded in the SC population response. Indeed, we have long appreciated that the population of SC neurons collectively codes for visual and motor behavior (Lee et al., 1988; McIlwain, 1991), but the actual mechanism by which the population activity is combined remains controversial. As suggested by computational models of perception that posit neural populations encode a probability distribution over sensory features (Ma et al., 2006), the population response in SC may encode the probability of a prioritized, including a remembered, location (Fecteau and Munoz, 2006; Kim and Basso, 2010). Bayesian decoding models of fMRI population responses have provided compelling evidence for probabilistic encoding in visual cortex (van Bergen et al., 2015), which could be extended to the SC. Overall, future research should address how cortical and subcortical brain areas differ in their support of WM. Perhaps, they utilize similar mechanisms, but differences may emerge due to their areal input-output connections, parameters that govern local circuit-level dynamics, or more broadly factors related to behavioral goals and task contexts (Sreenivasan and D’Esposito, 2019).

## Acknowledgements

We thank Jeffrey Kravitz for processing and analyzing the eye tracking data. We thank New York University’s Center for Brain Imaging for technical support. We thank Martin Paré for helpful discussions about the research study. This research was supported by the National Eye Institute (R01 EY-016407 and R01 EY-027925 to C.E. Curtis) and by an NYU Global Seed Grants for Collaborative Research (K.K. Sreenivasan and C.E. Curtis).

